# Intervertebral disc cell chondroptosis elicits neutrophil response in *Staphylococcus aureus* spondylodiscitis

**DOI:** 10.1101/2022.01.31.478414

**Authors:** Tiziano A. Schweizer, Federica Andreoni, Claudio Acevedo, Thomas C. Scheier, Irina Heggli, Ewerton Marques Maggio, Nadia Eberhard, Silvio D. Brugger, Stefan Dudli, Annelies S. Zinkernagel

## Abstract

**Objective:** To understand the pathophysiology of spondylodiscitis due to *Staphylococcus aureus*, an emerging infectious disease of the intervertebral disc (IVD) and vertebral body with a high complication rate, by combining clinical insights and experimental approaches.

**Design:** Clinical data and histological material of nine patients suffering from *S. aureus* spondylodiscitis were retrospectively collected at a single center. To mirror the clinical findings experimentally, we developed a novel porcine *ex vivo* model mimicking acute *S. aureus* spondylodiscitis and assessed the interaction between *S. aureus* and IVD cells within their native environment. In addition, the inflammatory features underlying this interaction were assessed in primary human IVD cells. Finally, mirroring the clinical findings, we assessed primary human neutrophils for their ability to respond to secreted inflammatory modulators of IVD cells upon *S. aureus* challenge.

**Results:** Acute *S. aureus* spondylodiscitis in patients was characterized by tissue necrosis and neutrophil infiltration. Additionally, the presence of empty IVD cells’ lacunae was observed. This was mirrored in the ex vivo porcine model, where *S. aureus* induced extensive IVD cell death, leading to empty lacunae. Concomitant engagement of the apoptotic and pyroptotic cell death pathways was observed in primary human IVD cells, resulting in cytokine release. Among the released cytokines, functionally intact neutrophil-priming as well as broad pro- and anti-inflammatory cytokines known for their involvement in IVD degeneration were found.

**Conclusions:** In patients as well as *ex vivo* in a novel porcine model, *S. aureus* spondylodiscitis infection caused IVD cell death, resulting in empty lacunae, which was accompanied by release of inflammation markers and recruitment of neutrophils. These findings offer valuable insights into the important role of inflammatory IVD cell death during the onset of spondylodiscitis and potential future therapeutic approaches.

## Introduction

The incidence of *Staphylococcus aureus* spondylodiscitis, an infection of the intervertebral disc (IVD) and adjacent vertebral bodies, has been steadily rising during the last two decades, classifying it as an emerging infectious disease (1,2). Spondylodiscitis is a severe disease, often resulting in IVD degeneration (3). This leads to drastic impairment of daily activities in the affected patients (4). *S. aureus* reaches the spine via hematogenous seeding during bacteremia, per continuitatem by spread from local soft tissue infections or by direct injection, such as unintentional inoculation during medical procedures (5-7). Due to the absence of immune cells and limited vascularization in the IVD, which impairs the penetration of the antibiotics into the IVD, prolonged antibiotic treatment for a minimum of six weeks is required in order to treat the bacterial infection (8,9). However, the pathophysiology of *S. aureus* spondylodiscitis resulting in IVD destruction and therefore potential crucial insights, which would allow to improve treatment, remains still vastly unknown, among others due to the lack of adequate models.

Since animal models of spondylodiscitis are work-intensive and only allow low throughput and read-outs, we aimed at establishing a porcine *ex vivo* model to assess the initial interaction between *S. aureus* and IVD cells in their native environment. Porcine models are more frequently used as preclinical models in infectious diseases, since their anatomical, physiological and immunological response properties are close to those of humans (10-12). Furthermore, pigs, as well as humans, are frequently colonized with *Staphylococcus* spp. and are among the livestock animals with reported cases of spondylodiscitis, caused predominantly by *Streptococcus* and *Staphylococcus* spp. (13).

The human adult IVD contains only two distinct types of IVD cells, nucleus pulposus cells found in the center and annulus fibrosus cells found in the outer margins of the IVD (14). IVD cells possess phagocytic capacity, allowing them to remove dead IVD cells, but also, as recently shown, to phagocytose live *S. aureus* via the Toll-like receptor 2 (TLR2) pathway (15–17). However, the fate of IVD cells after encountering *S. aureus* still remains unknown. In contrast to the fulminant infection caused by *S. aureus*, the Gram-positive commensal *Cutibacterium acnes* can cause low-grade chronic IVD infections, during which caspase-mediated IVD cell death following TLR2 recognition was reported (18), suggesting a key role for regulated cell death in spondylodiscitis initiation.

Regulated IVD cell death has been extensively studied in connection to its potential role in IVD degeneration. Studies showed that in sterile IVD inflammation, IVD cells frequently undergo regulated cell death morphologically similar to chondroptosis (19,20). Chondroptosis was first described for articular chondrocytes in the growth plate of rabbit femurs (21,22) and considered to be an apoptotic-derived regulated cell death type, characterized by the activation of apoptotic caspases and formation of autophagic vacuoles to sequester and degrade cellular material (23). Furthermore, IVD cells have the ability to express and secrete a wide array of cytokines in response to various environmental stimuli (24–27). Cytokine secretion might be enhanced during chondroptosis of IVD cells and potentially cause a strong local immune stimulation (28,29).

In the present study, we aimed at characterizing the pathophysiology of *S. aureus* spondylodiscitis. We used a multipronged approach, starting with dissecting the clinical and histological presentation of *S. aureus* spondylodiscitis, translating this into a newly established porcine *ex vivo* model of spondylodiscitis and finally characterizing molecular changes occurring in primary human IVD cells upon *S. aureus* challenge as well as the neutrophil-recruitment and -activation potential of IVD cells. The findings of our study deliver important insights into the pathophysiology of *S. aureus* spondylodiscitis and into potential alternative treatment approaches.

## Results

### Acute *S. aureus* spondylodiscitis is characterized by inflamed and necrotic tissue with signs of IVD cell death and neutrophil infiltration

The medical documentation of a case series of nine hospitalized spondylodiscitis cases at the University Hospital Zurich, Switzerland, with a positive *S. aureus* blood culture was reviewed (**Table 1**). The median age was 58 years. Four patients had at least one predisposing factor of contracting infectious diseases and five patients had no identifiable comorbidities. The female to male ratio was 5:4. All patients showed elevated c-reactive protein (CRP) and white blood cell count (WBC) (**Table 2**). The length of hospital stay (LOS) ranged from ten to 68 days, with a median LOS of 21 days and a 30 day survival rate of 77.8%. Out of the nine patients, six showed radiological signs of intervertebral disc degeneration with edema (**Figure 1, A and B**). Histological IVD material was only available for patient 3 (acute state) and 6 (chronic state). In patient 3, Brown-Brenn staining identified Gram-positive cocci (**Figure 1, C and D**). Hematoxylin & Eosin staining showed strongly inflamed and necrotic tissue, characterized by the presence of empty lacunae, an indication for IVD cell death (**Figure 1, E and F**). Additionally, neutrophil infiltration in the IVD tissue was observed (**Figure 1, G and H**). In contrast, in the histology sample from patient 6, the inflammatory infiltrate was mostly composed of lymphocytes and no bacteria were observed (**Figure S1**).

**Table 1.**
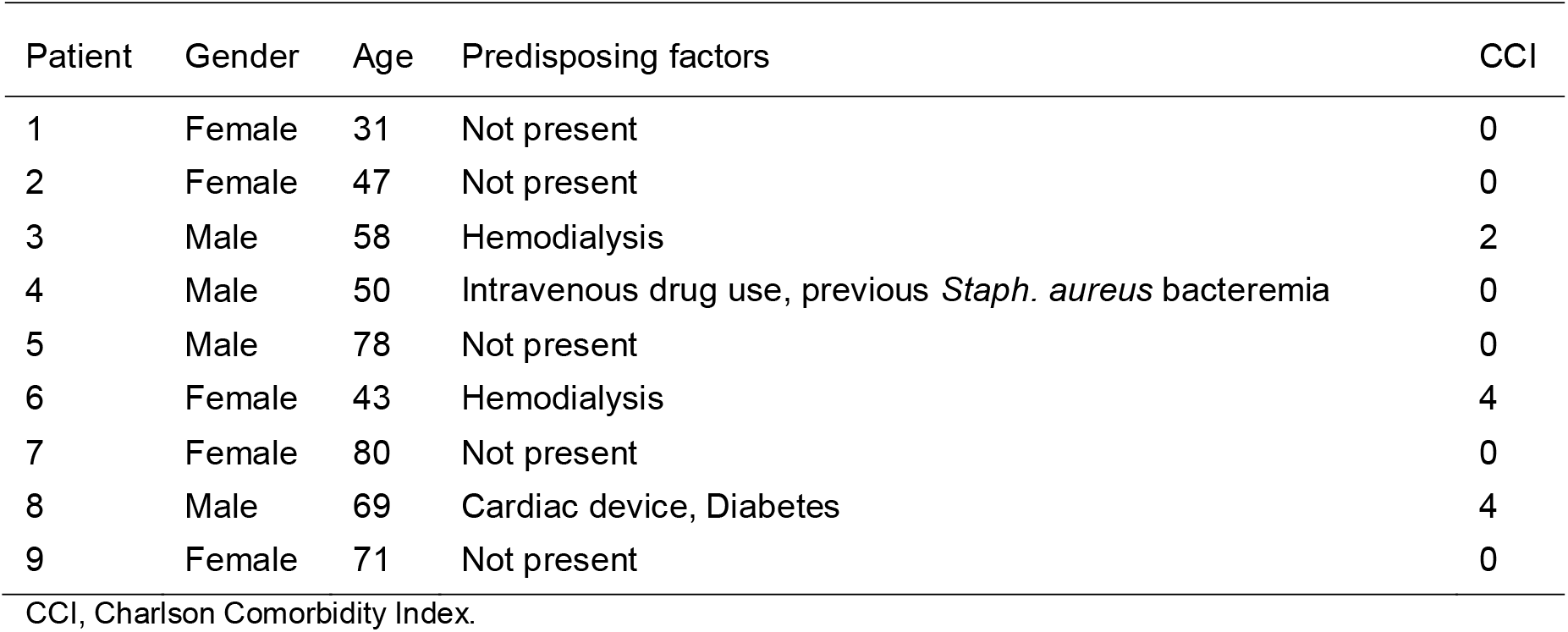
Demographic overview of spondylodiscitis patients.

| Patient | Gender | Age | Predisposing factors | CCI |
| --- | --- | --- | --- | --- |
| 1 | Female | 31 | Not present | 0 |
| 2 | Female | 47 | Not present | 0 |
| 3 | Male | 58 | Hemodialysis | 2 |
| 4 | Male | 50 | Intravenous drug use, previous *Staph. aureus* bacteremia | 0 |
| 5 | Male | 78 | Not present | 0 |
| 6 | Female | 43 | Hemodialysis | 4 |
| 7 | Female | 80 | Not present | 0 |
| 8 | Male | 69 | Cardiac device, Diabetes | 4 |
| 9 | Female | 71 | Not present | 0 |

CCI, Charlson Comorbidity Index.

**Table 2.** Clinical characteristics and laboratory parameters of spondylodiscitis patients at admission.

| Patient | Site of Acquisition | Source of Infection | Bacteremia | Localization | Radiological Signs of Degeneration | LOS [days] | 30 Day Survival | CRP [mg/l] | WBC [G/l] |
| --- | --- | --- | --- | --- | --- | --- | --- | --- | --- |
| 1 | CA | Unknown | Yes | Th10 - Th11 | No | 19 | Yes | 167 | 14.83 |
| 2 | CA | Unknown | Yes | L2 - L3 | Yes | 22 | Yes | 403 | 21.37 |
| 3 | HC | Hemodialysis Catheter | Yes | Th11 - Th12 | Yes | 25 | Yes | 88 | 9.32 |
| 4 | CA | Unknown | No | L1 - L3 | Yes | 10 | Yes | 196 | 7.28 |
| 5 | CA | Unknown | Yes | C6 - C7 | Yes | 15 | Yes | 487 | 15.3 |
| 6 | HC | Unknown | Yes | Th8 - Th9 | No | 23 | Yes | 86 | 8.24 |
| 7 | CA | Unknown | Yes | Th10 - Th11 | Yes | 21 | No | n/a | n/a |
| 8 | CA | Device-related infection | Yes | L2 - L4 | Yes | 68 | Yes | 339 | 15.07 |
| 9 | CA | Soft Tissue Infection | Yes | L1 - L4 | n/a | 20 | No | 144 | 31.2 |

LOS, length of stay; CRP, c-reactive protein; WBC, white blood cell count; CA, community acquired; HC, healthcare related; Th, thoracic, L, lumbar, C, cervical;

**Figure 1.**
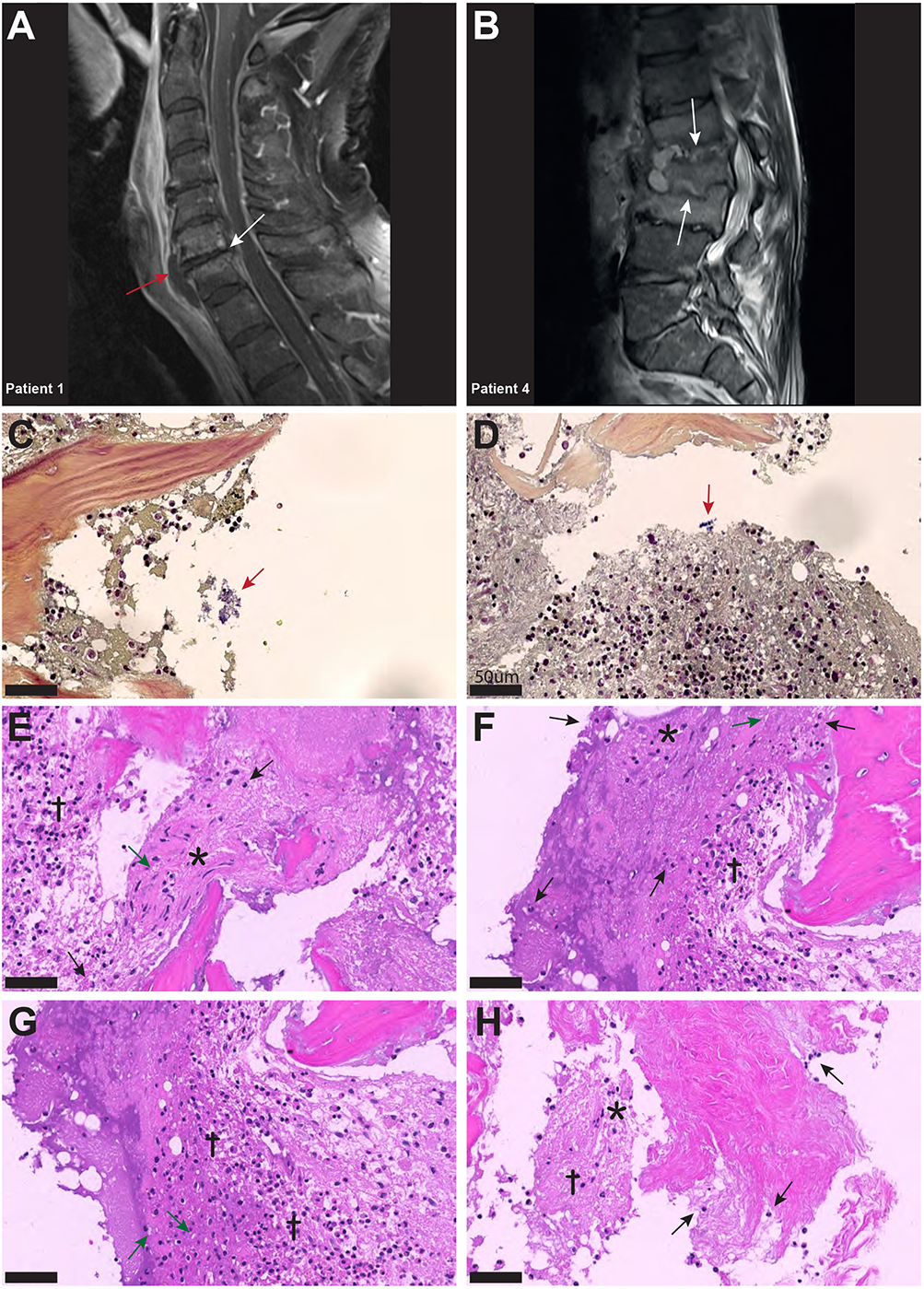
Inflammatory infiltrates with presence of empty IVD cells’ lacunae and neutrophils in IVD during acute *S. aureus* spondylodiscitis. (**A and B**) T1 weighted MRI scans showing inflammatory infiltrates into and degenerative changes of the IVD in patient 1 and 4. White arrows indicate localization of spondylodiscitis, red arrow indicates paravertebral abscess. (**C and D**) BB staining showing the presence of Gram positive cocci (red arrow) in the IVD of patient 3. (**D-H**) H&E staining showing strongly inflamed and necrotic IVD tissue (cross), clusters of IVD cells (asterisk), the presence of empty IVD cells’ lacunae (green arrow) and the infiltration of immune cells, mostly neutrophils, into the IVD (black arrow). Scale bars indicate 50 μm. BB, Brown-Brenn; H&E, Hematoxylin & Eosin, IVD, intervertebral disc.

### *S. aureus* challenge induces chondroptosis of IVD cells in the *ex vivo* porcine spondylodiscitis model

The IVDs were removed from the spines, followed by preparation of annulus fibrosus IVD punches with a biopsy punch (**Figure S2, A and B**). The IVD punches were challenged with a patient-derived *S. aureus* strain, to assess whether *S. aureus* could grow and persist in the IVD tissue environment. Transmission electron microscopy (TEM) and histology confirmed the presence of *S. aureus* deeply embedded within the IVD punch (**Figure 2, A and F**). *S. aureus* grew within the IVD environment for up to 48h (**Figure 2B**). When comparing different laboratory and clinical strains (**Table S1**), differences in the potential of initial IVD attachment and growth were observed (**Figure 2C**). Of note, staphylococci were also identified within (**Figure 2D**) and close to lysed IVD cells, surrounded by cellular matrix (**Figure 2E**), an indication for *S. aureus* cytotoxcity towards IVD cells.

**Figure 2.**
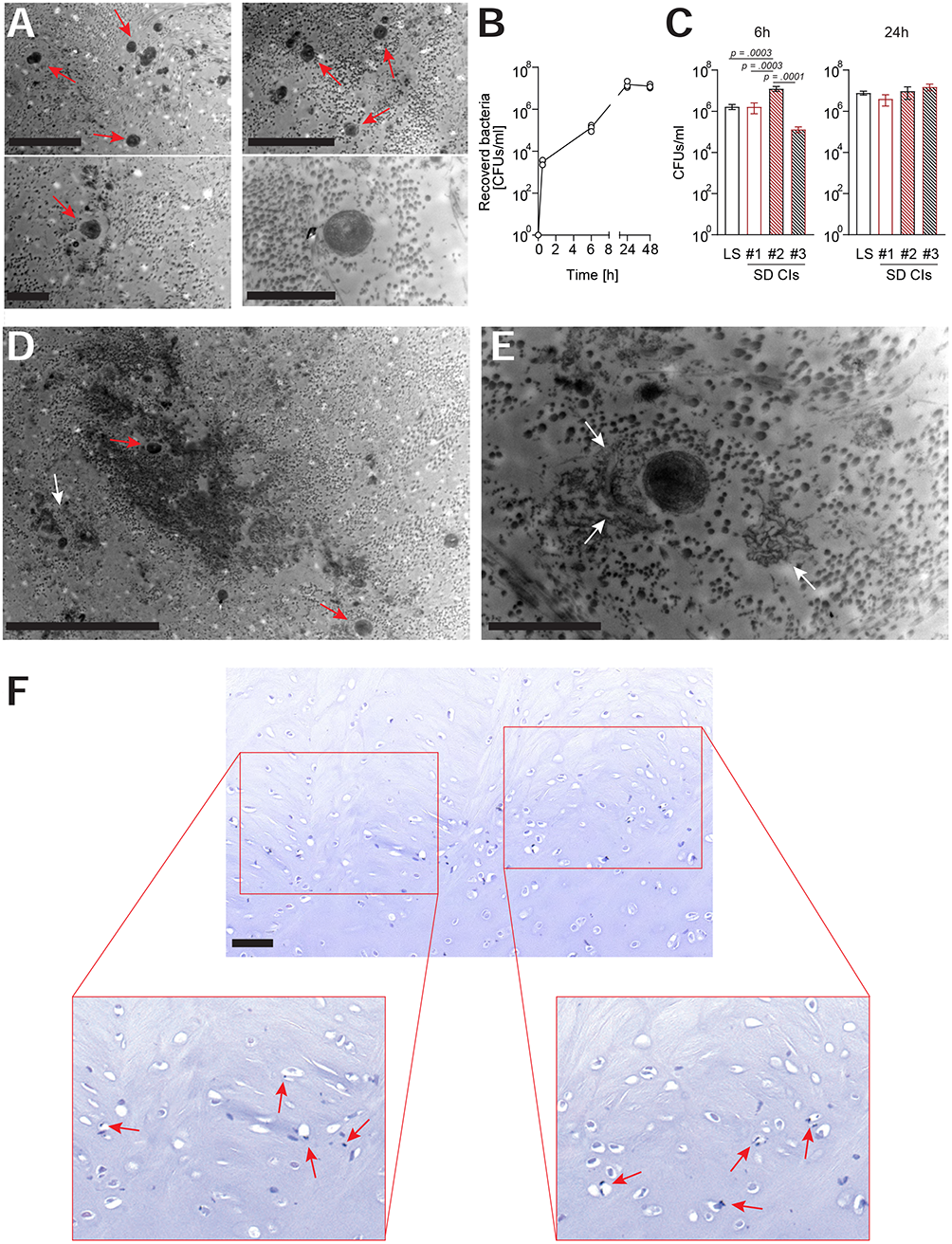
*S. aureus* rapidly grows and maintains within a novel porcine *ex vivo* spondylodiscitis model. (**A**) TEM showing single cells and clusters of Staphyloocci deep within the porcine IVD. (**B**) Growth curve of a *S. aureus* laboratory strain within the IVD punches over 48 h. (**C**) Comparison of the ability of different *S. aureus* spondylodiscitis (SD) isolates to colonize and grow within the IVD punches. (**D and E**) Overview of multiple staphylococci (red arrow) within and around a lysed IVD cell (D) and close-up of single *S. aureus* surrounded by cellular material (white arrow) (E). (**F**) Gram staining of *S. aureus* challenged IVD punch, showing the presence of *S. aureus* deep within the IVD punch (red arrows). Data are presented as mean ± standard deviation from three biological replicates. Statistical analysis was done by one-way ANOVA and Turkey’s multiple comparisons test. Scale bars indicate 5 μm and 2 μm (A, top and bottom panels, respectively), 10 μm (D), 2 μm (E) and 50 μm (F). TEM, transmission electron microscopy; IVD, intervertebral disc.

Histological analysis showed an increase in the presence of empty lacunae, an indication for IVD cell death (22), in punches challenged with *S. aureus* (**Figure 3, A and B**). In order to understand whether IVD cells underwent regulated cell death, IVD cells isolated from IVD punches challenged or not with *S. aureus* were assessed by flow cytometry using the Annexin V/7AAD staining method (**Figure S3A**). The isolated IVD cells were very pure and showed no signs of contamination by either immune or endothelial cells (**Figure S3B**). IVD cells isolated from *S. aureus* challenged IVD punches showed a significantly lower proportion of viable cells and a higher proportion of cells undergoing regulated cell death as compared to IVD cells isolated from uninfected IVD punches (**Figure 3, C and D**). Morphological analysis by TEM revealed the loss of the usually elongated fibroblastic phenotype of annulus fibrosus IVD cells, when challenged with *S. aureus* (**Figure 3E and S3C**). Nuclei showed signs of local degradation but no dispersion in the cellular lumen. Furthermore, membrane integrity seemed to be lost and many vacuoles were observed (**Figure 3E and F**). Additionally, cellular debris was located at the border of the lacunae (**Figure 3G**).

**Figure 3.**
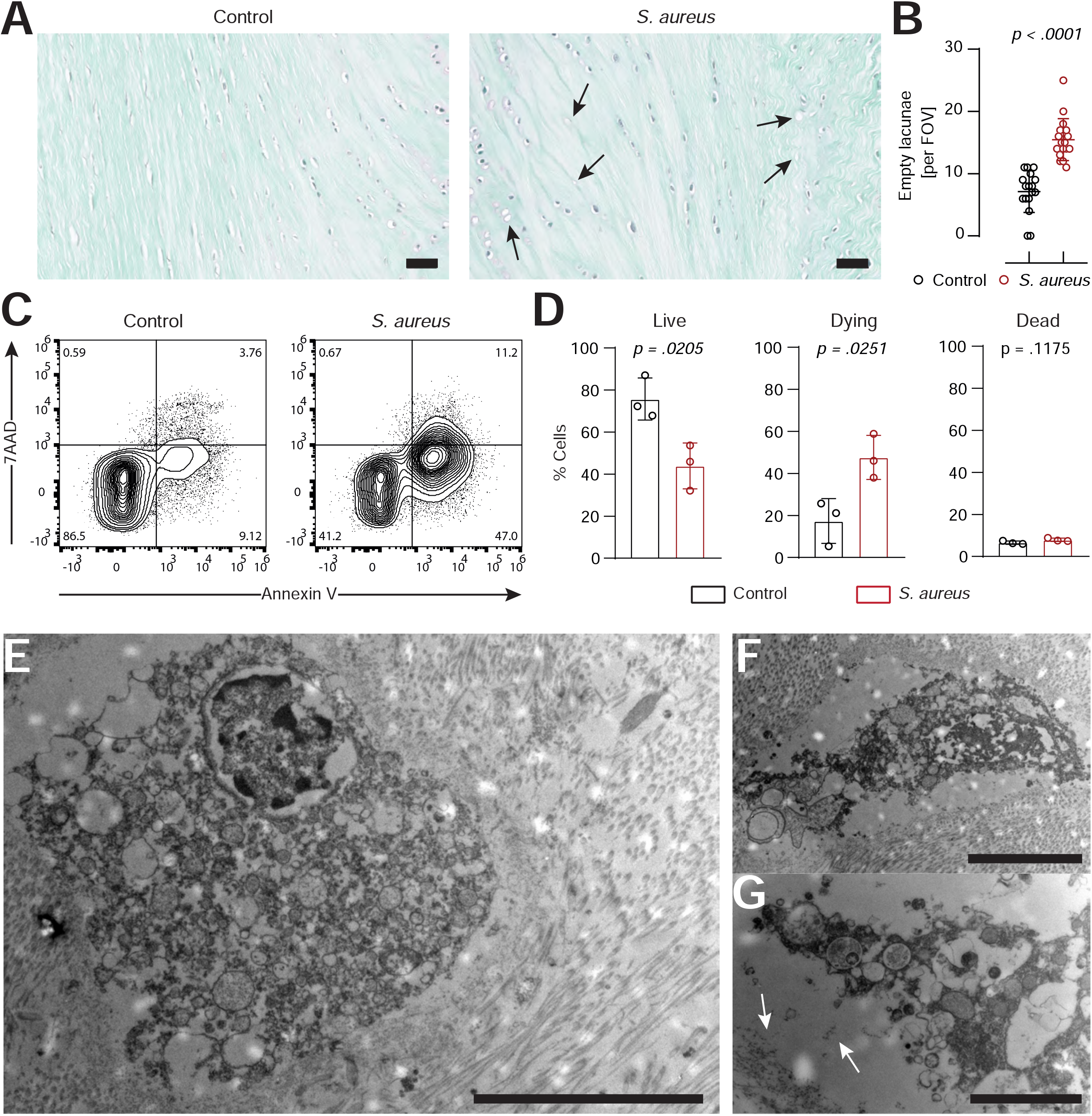
IVD cells in the porcine ex *vivo* model undergo extensive chondroptosis upon *S. aureus* challenge. (**A and B**) Safranin-o/fast-green staining of histological slides (A) and quantification of empty lacunae in IVD punches either unchallenged or challenged with *S. aureus* for 24h. Dark arrows indicate empty lacunae. In total, 17 randomly selected FOVs from six different IVD punches per group were chosen to evaluate the presence of empty lacunae. (**C and D**) Representative flow cytometry plot of isolated IVD cells from either unchallenged or *S. aureus* challenged IVD punches and stained with AnnexinV/7AAD (C) and quantification of live (Annexin V^-^, 7AAD^-^), dying (Annexin V^+^, 7AAD^-^) and dead (Annexin V^+^, 7AAD^+^) IVD cells (D). (**E-G**) TEM of IVD cells found within S. aureus challenged IVD punches. White arrows in G indicate the deposition of cellular material to the edge of the lacuna. In D, data are presented as mean ± standard deviation from three biological replicates, where each dot represents one biological replicate. Statistical analysis was done by paired t-test. Scale bars indicate 100 μm (A), 10 μm (E), 5 μm (F) and 2 μm (G). IVD, intervertebral disc; FOV, field of view; TEM, transmission electron microscopy.

### *S. aureus-*induced chondroptosis in human IVD cells is linked to apoptotic and pyroptotic caspases

To better understand the molecular mechanisms underlying the IVD cells *S. aureus* interaction, we assessed the response of primary human annulus fibrosus IVD cells to *S. aureus* challenge. Isolated primary human IVD cells (**Figure S4, A and B**) efficiently phagocytosed, but did not eradicate *S. aureus* (**Figure S4, C and D**). Challenge with either *S. aureus* or *S. aureu*s conditioned-medium caused IVD cells to undergo regulated cell death (**Figure 4A, Figure S4E**). Visual assessment by confocal laser scanning microscopy revealed altered IVD cells morphology (**Figure 4, B and C**) resembling the hallmarks of chondroptosis, such as membrane blebbing and the presence of vacuoles with intact nuclei, when challenged with *S. aureus*, which was not observed in unchallenged IVD cells (**Figure 4, D and E**). Next, we assessed whether caspases were involved in *S. aureus-*induced chondroptosis. Indeed, we found elevated activity of the effector caspase-3/7 as well as the initiator caspase-8 (**Figure 4F**), whereas no changes in caspase-9 activity were observed. Caspase-8 inhibition led to significantly enhanced survival, while caspase-9 inhibition did not block cell death induction, further confirming our findings (**Figure 4G**). Since we previously observed that porcine IVD cells presented with loss of membrane integrity, we also investigated whether chondroptosis upon *S. aureus* challenge involved changes to the membrane permeability of human IVD cells. Although no significant differences in cellular ATP levels were found, *S. aureus* challenge led to an increased loss of membrane integrity (**Figure 4H**). The observed vacuoles might represent lysosomal exocytosis, i.e. for transport of cytokines to the extracellular space. Therefore, we assessed CD107a expression on IVD cells as a marker for lysosomal exocytosis. We found significantly higher expression of CD107a on IVD cells challenged with *S. aureus* as compared to uninfected cells (**Figure 4I**). Furthermore, we observed increased caspase-1 activity in infected IVD cells (**Figure 4I**), in line with IL-1β and IL-18 secretion (**Figure 4J**). Moreover, pan-caspase inhibition showed a significantly higher effect on IVD cells survival than either caspase-1 or caspase-8 inhibition alone (**Figure S4F**). Finally, immunohistochemistry (IHC) of patient histology showed positive staining for cleaved caspase-3 and caspase-1 in IVD cells during *S. aureus* spondylodiscitis (**Figure 4K**).

**Figure 4.**
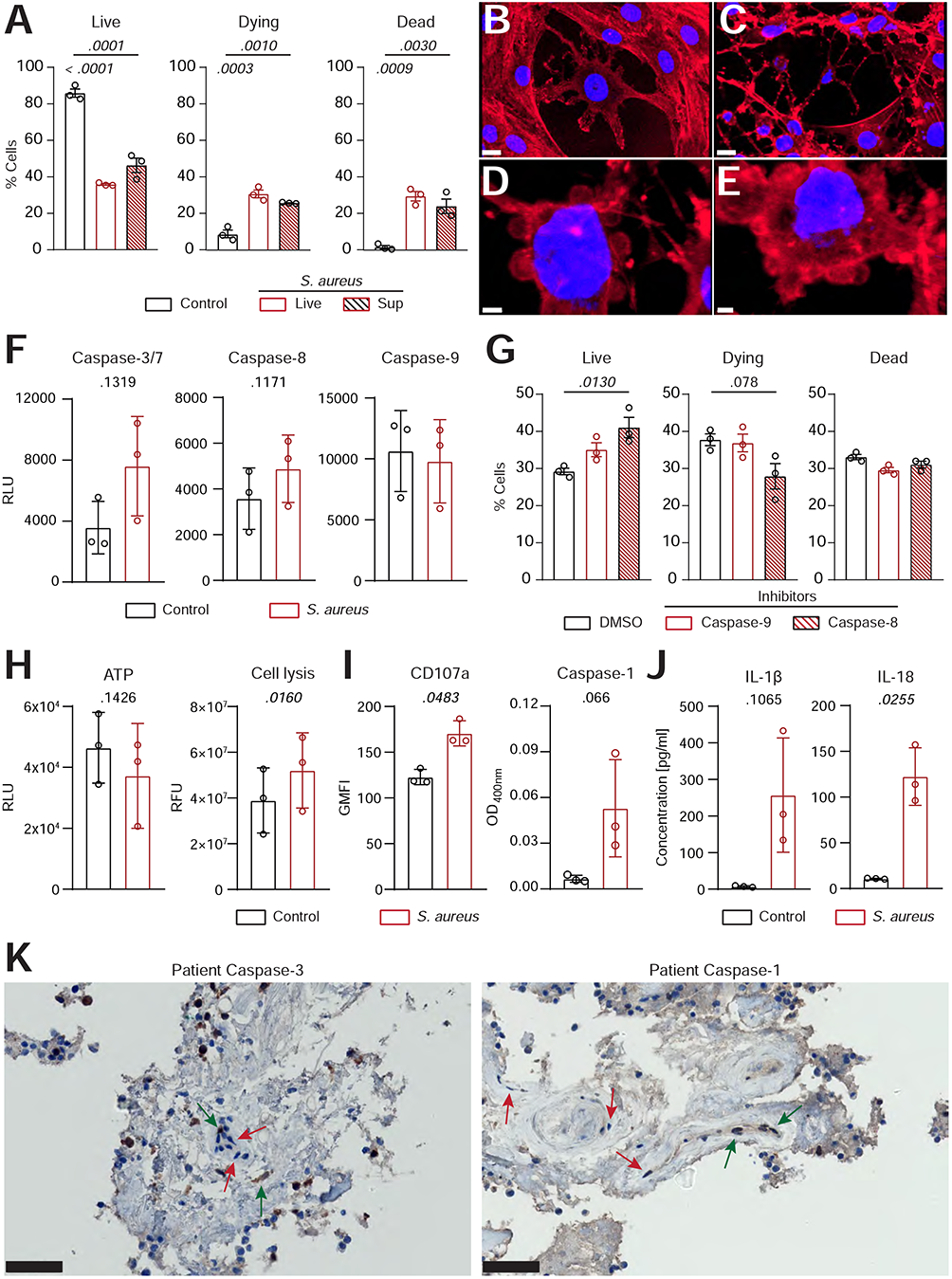
Human primary IVD cells undergo chondroptosis linked to concomitant activation of the apoptotic and pyroptotic pathway upon *S. aureus* challenge. (**A**) Isolated human primary IVD cells were infected with the clinical *S. aureus* isolate from patient 7 at MOI 10 or *S. aureus* supernatant for 24h and viability was assessed by flow cytometry with Annexin V/7AAD, quantifying live (Annexin V^-^, 7AAD^-^), dying (Annexin V^+^, 7AAD^-^) and dead (Annexin V^+^, 7AAD^+^) IVD cells. (**B-E**) Representative CLSM images showing an overview of unchallenged IVD cells (B) and IVD cells challenged with live *S. aureus* (C-E). Cells were stained with Hoechst (blue, nucleus) and Rhodamine-phalloidin (red, actin cytoskeleton). (**F**) Analysis of caspase-3/7, caspase-8 and caspase-9 activity in unchallenged or challenged IVD cells. (**G**) Flow cytometry analysis of challenged IVD cells in the presence or absence of caspase-8 or caspase-9 inhibitors (50 μM), quantifying live, dying and dead cells. (**H and I**) Analysis of ATP levels, cell membrane integrity and marker for lysosomal exocytosis (CD107a) (H) and caspase-1 activity (I) in unchallenged or challenged IVD cells. (**J**) Luminex-based analysis of IL-1β and IL-18 secretion into supernatant by unchallenged or challenged IVD cells. (**K**) IHC of cleaved caspase-3 and caspase-1 in the IVD of patient 3, showing stained (green arrow) and unstained (red arrow) IVD cells. Data are presented as mean ± standard deviation from three biological replicates, where each dot represents one biological replicate. Statistical analysis was done by one-way ANOVA and Turkey’s multiple comparisons test or paired t-test. Scale bars indicate 100 μm (D) and 50 μm (K). IVD, intervertebral disc; MOI, multiplicity of infection; CLSM, confocal laser scanning microscopy; IHC, immunohistochemistry.

### IVD cells secrete functional neutrophil-priming cytokines upon *S. aureus-*induced chondroptosis

Since challenged IVD cells secreted the pro-inflammatory cytokines IL-1β and IL-18, we investigated whether they also secreted neutrophil-targeting cytokines. *S. aureus* challenged IVD cells secreted significantly elevated levels of IL-8, G-CSF, CXCL1, CXCL2 and CXCL12 (**Figure 5A**). Apart from IL-8 and CXCL12, the levels of secreted cytokines where higher at later timepoints of infection. To assess whether the secreted cytokines were functional, isolated human neutrophils were stimulated with the filtered supernatant of infected or uninfected IVD cells. Neutrophils stimulated with supernatant from infected IVD cells showed significantly higher expression of CD66b and CD15 as compared to neutrophils stimulated with supernatant from uninfected IVD cells or medium only (**Figure 5C, Figure S5, A-C**). Furthermore, the corresponding receptors, CXCR1 for IL-8, G-CSF and CXCL1, CXCR2 for IL-8 and CXCL2 and CXCR4 for CXCL12, showed decreased expression upon stimulation with supernatant from challenged IVD cells, pointing towards active engagement of these receptors (30). This was only observed upon stimulation with supernatant from IVD cells at late infection stages, while an early infection stage supernatant did not yield the same effect, making bacterial toxins as a possible bias unlikely (**Figure S5D**). We found that significantly more neutrophils migrated towards supernatants from infected IVD cells as compared to supernatants from uninfected IVD cells or medium alone (**Figure 5E**) and they also produced higher reactive oxygen species (ROS) levels (**Figure 5F**). Finally, we also explored the potential of IVD cells to secrete additional cytokines, involved in a more comprehensive immune response. IVD cells secreted a broad range of cytokines upon *S. aureus* challenge, such as the general inflammation and damage markers IL-1α, IL-6, TNF-α and TNF-β, the monocyte stimulating cytokines GM-CSF, M-CSF, MIP-1α and MIP-1β, the T cell stimulating cytokines IL-2, IL-12p70 and IL-17A, as well as the anti-inflammatory cytokines IL-4 and IL-10 (**Figure 5G**).

**Figure 5.**
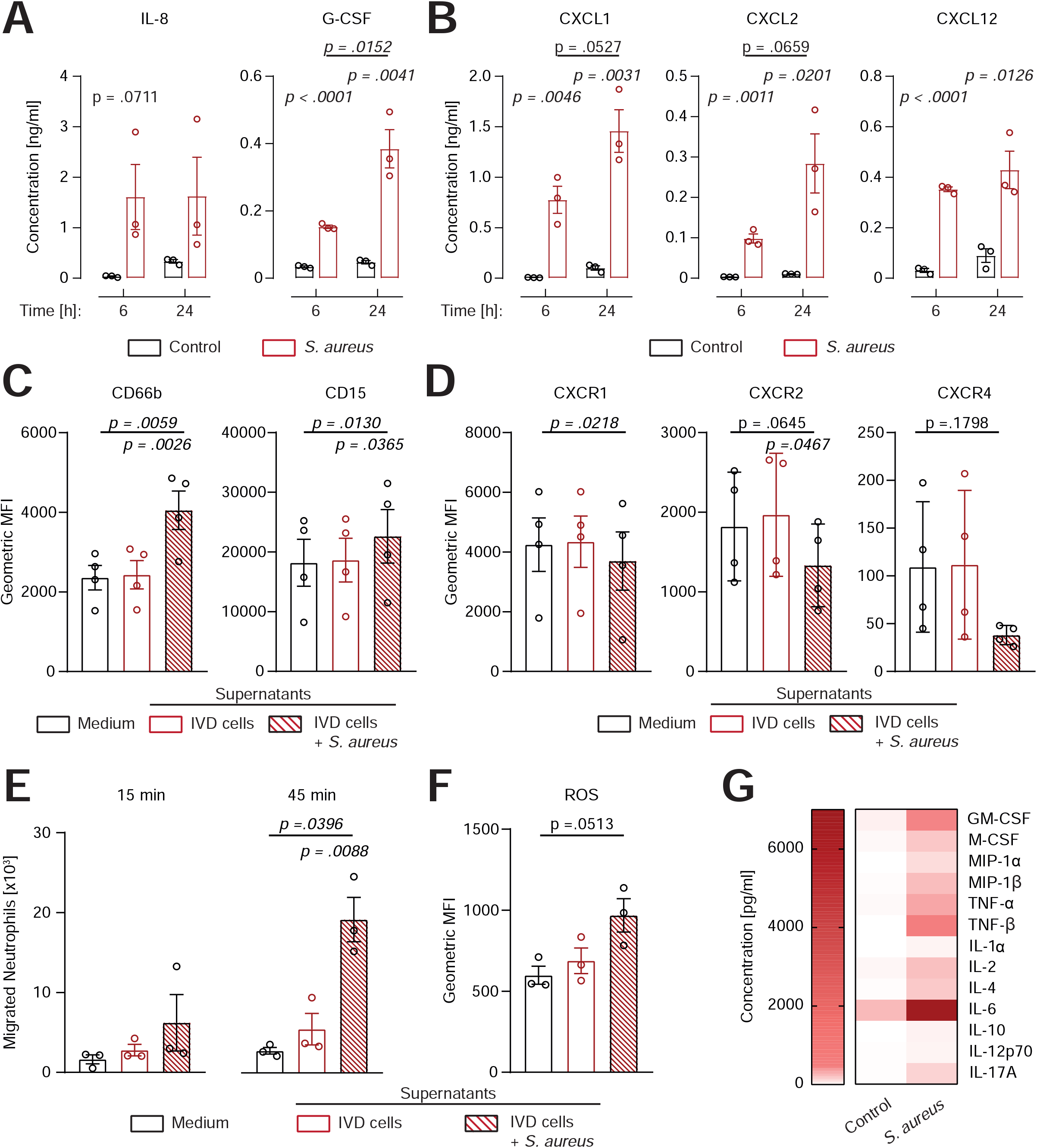
Human IVD cells secrete a broad range of cytokines upon *S. aureus* challenge, including functional neutrophil-priming cytokines. (**A and B**) Luminex-based analysis of IL-8, G-CSF (A) as well as CXCL1, CXCL2 and CXCL12 (B) secretion into supernatant by unchallenged or *S. aureus-*challenged IVD cells. (**C and D**) Activation and maturation markers expression (C) and chemokine receptor expression (D) on primary human neutrophils stimulated with medium only or cell culture supernatant from either unchallenged or challenged IVD cells. (**E**) Migration assays of primary human neutrophils towards medium only or cell culture supernatant from either unchallenged or challenged IVD cells. (**F**) Assessment of ROS production in primary human neutrophils stimulated with medium only or cell culture supernatant from either unchallenged or challenged IVD cells. (**G**) Luminex-based analysis of broad pro-and anti-inflammatory cytokine secretion into supernatant by unchallenged or challenged IVD cells. Data are presented as mean ± standard deviation from at least three biological replicates, each dot represents one biological replicate. Statistical analysis was done by two-way ANOVA and Sidak’s multiple comparisons test or one-way ANOVA and Turkey’s multiple comparisons test. IVD, intervertebral disc; ROS, reactive oxygen species.

## Discussion

In patients suffering from *S. aureus* spondylodiscitis, we identified the presence of empty lacunae, an indication for IVD cell death, and the influx of neutrophils into the IVD. These findings were mirrored experimentally, by establishing a novel porcine spondylodiscitis model and by using primary human IVD cells. We confirmed that IVD cells underwent chondroptosis linked to a strong immune-stimulating cytokine secretion profile, especially directed towards the activation and recruitment of neutrophils. The strength of our study lies in the combination of clinical findings and novel spondylodiscitis-specific *ex vivo* models.

In our case series analysis of *S. aureus* spondylodiscitis cases, we observed increased inflammation as well as IVD degeneration, as reported previously for spondylodiscitis (31-33). For the first time, we show Gram-positive cocci and the presence of empty IVD cells’ lacunae, which was not documented in the literature so far. This clinical observation shows interaction between IVD cells and Gram-positive cocci in the human IVD.

In order to investigate these clinical findings in detail, we developed a novel porcine *ex vivo* spondylodiscitis model. Of note, IVD cells found in the porcine nucleus pulposus might still resemble a stem cell like population, which is only found in humans during their infancy (34). Therefore, we used IVD punches prepared exclusively from the annulus fibrosus of porcine IVDs. *S. aureus* was able to grow to high densities and persist within the IVD environment, as previously reported for other bacteria (18,25,35). Challenge with *S. aureus* led to accumulation of empty lacunae within the IVD, an indication for regulated IVD cell death (23,36), confirmed by Annexin V staining. However, since Annexin V staining is just an indicator for any type of regulated cell death, it is crucial to further determine the involved regulated cell death pathways (37). Since chondroptosis is regarded as the *de facto* regulated IVD cell death (20,23), it is not surprising that IVD cells underwent chondroptosis upon *S. aureus* challenge, as corroborated by TEM, especially since it was previously observed that IVD cells underwent caspase-mediated regulated cell death upon *C. acnes* challenge (18). One can assume that chondroptosis might be a conserved response of IVD cells towards bacterial encounter.

Chondroptosis was also observed in primary human IVD cells upon *S. aureus* challenge. Importantly, supernatants of staphylococcal cultures were sufficient to induce chondroptosis, potentially attributable to the presence of secreted toxins. Among them, the staphylococcal α-toxin, a potent inducer of cell death in many different cell types, including articular chondrocytes, might play a key role in inducing chondroptosis in IVD cells (38–40), apart from TLR2 engagement by bacterial associated molecular patterns (18).

We observed that chondroptosis of human IVD cells upon *S. aureus* challenge was linked to increased caspase-3/7 and caspase-8 activity, but not caspase-9 activity. Caspase-3/7 activation was previously shown in IVD cells in IVDD (41–44). Both caspase-8 and caspase-9 activation were shown to occur in IVD cells during sterile IVD inflammation (45,46). However, in our study, IVD cells only showed increased caspase-8 but not caspase-9 activity. Furthermore, we found that chondroptosis upon *S. aureus* challenge involved cell membrane integrity loss and lysosome formation. Lysosomal exocytosis is a known mechanism of pro-inflammatory cytokine secretion, best known for IL-1β and IL-18 (28,29). Both IL-1β and IL-18 are stored within the cell in an inactive pro-form, which needs to be cleaved and activated by proteolytic enzymes, such as caspase-1 (47). This suggests that also caspase-1, which indeed showed increased activity, played an important role in chondroptosis upon *S. aureus* challenge. Our findings of IL-1β and IL-18 secretion together with elevated caspase-1 activity in chondroptotic IVD cells were in line with findings of the engagement of the NLRP3/caspase-1 axis in IVD cells with IL-1β secretion in IVDD (24,48–51). The IHC findings of the patient spondylodiscitis, although depicting a situation during the course of disease rather than at the onset, further corroborated the experimental findings. It was recently shown that regulated cell death pathways, i.e. apoptosis, necroptosis and pyroptosis, referred to as PANoptosis, are intermittently linked and regulated together (52). Given our findings of concomitant activation of apoptotic and pyroptotic caspases, chondroptosis seems to follow a conserved pattern of regulated cell death activation, relying on multiple initiation and execution platforms.

Previous research showed that IVD cells express and secrete neutrophil-priming cytokines upon IL-1β stimulation (53,54). We further corroborated these findings in the context of *S. aureus* challenge of IVD cells and showed that the secreted cytokines were functionally intact, leading to neutrophil priming. However, whether neutrophil recruitment and activation in *S. aureus* spondylodiscitis is beneficial to clear the infection or detrimental, resulting in damaged tissue, still needs to be elucidated, since extensive ROS release causes tissue injury (55). Nevertheless, these findings add to the growing body of evidence that IVD cells are potent immune cells’ recruiter, as previously shown for T cells (26). Apart from neutrophil or T cell-specific cytokines, various studies showed that the pro-inflammatory cytokines TNF-α, TNF-β, IL-1α, IL-1β, IL-6 and IL-17A, also secreted after *S. aureus* challenge in our study, cause IVDD by inducing regulated cell death, stimulating matrix degradation, chemokine secretion and changes in IVD cells’ phenotype (56). Our findings suggest that acute *S. aureus* challenge of IVD cells induces a similar phenotype as found in low-grade chronic IVDD. These findings have important implications and suggest that future therapeutic approaches should explore the potential impact of employing cytokine signaling interference or cell death inhibitors as complementary therapy to antibiotics in *S. aureus* spondylodiscitis.

## Material and Methods

### Ethical requirements

Use of patient-derived IVD material (Business Administration System for Ethics Committees [BASEC] No. 2018-01486) as well as clinical data of spondylodiscitis patients ([BASEC] No. 2016-00145, 2017-02225 and 2017-01140) and isolation of human neutrophils ([BASEC] No. 2019– 01735) was done in accordance with the Helsinki Declaration and approved by the Cantonal Ethical Research Committee Zurich. The medical documentation of nine patients suffering from spondylodiscitis and treated at the University Hospital of Zurich, Switzerland, between 2011 and 2020 was reviewed. The diagnosis of spondylodiscitis was based on microbiological, clinical and radiological parameters as well as laboratory tests. Porcine spines were obtained from freshly euthanized animals either from a local butcher (Metzgerei Angst, Zurich) or from the Center for Surgical Research at the University Hospital Zurich.

### Histology

Patient biopsies and porcine spine punches were fixed in 4% buffered formalin and paraffin embedded. Sections (2 μm) were stained with hematoxylin and eosin (H&E), Brown-Brenn (BB), Safranin-o & Fast-green or Gram-staining. For IHC, formalin-fixed paraffin-embedded tissue sections were pre-treated with the BOND Epitope Retrieval Solution 2 (Leica Biosystem) at 100°C for 30 min. They were stained with anti-Cleaved Caspase-3 antibody (polyclonal, ab2302, abcam) and Caspase-1 antibody (polyclonal, ab62698, abcam) for 30 min. For detection, the slides were stained with the Bond Polymer Refine Detection HRP Kit (Leica Biosystem), according to the manufacturer’s instruction and counterstained with haematoxylin. Whole-slide scanning and photomicrography were performed with a 108NanoZoomer 2.0-HT digital slide Scanner (Hamamatsu, Houston, TX).

### Bacterial strains and growth conditions

*S. aureus* JE2 USA300 (NARSA) and the clinical isolates were maintained on blood agar plates (Columbia agar +5% sheep blood, BD) and grown in Tryptic Soy Broth (TSB, BD) at 37°C and 220 rpm for 16 h. Cultures were diluted in fresh TSB and grown to exponential phase for the challenge of IVD cells. For supernatants challenge, overnight cultures were diluted in fresh DMEM/F12 (Gibco) + 10% FCS and grown to exponential phase (approx. 3h). The cultures were filter sterilized with a 0.22 μm filter and used in a ¼ dilution.

### Preparation and infection of porcine IVD punches

Porcine spines were cleaned with 70% EtOH wipes. The IVDs were removed along the endplates on both side and annulus fibrosus IVD punches were prepared with a 5 mm biopsy punch (Kai Medical). IVD punches were dissected longitudinally in the center and placed with the IVD side facing upwards in 96-well plates in DMEM/F12 + 10% FCS. IVD punches were infected with bacteria at an inoculum of 4×10^5^ colony forming units (CFUs) and incubated at 37°C + 5% CO_2_.

### Assessment of bacterial growth within IVD punches

At indicated timepoints, the medium was removed and the IVD punches were transferred to new wells. They were washed twice with DPBS and sonicated for 3 min to remove adhering bacteria. Next, they were placed into tubes with metallic beads and homogenized in a TissueLyser (Qiagen) at 30Hz for 10 min. The tubes were centrifuged at 1200 rpm for 3 min after which the supernatant was collected into fresh tubes and centrifuged at 14000rpm for 3 min. The resulting pellet was lysed with water, serially diluted and spotted on TSB agar (TSA) plates.

### Assessment of cell death of IVD cells within IVD punches

Washed IVD punches were fragmented into four pieces and enzymatically digested with enzymatic digestion buffer consisting of DMEM/F12 (Gibco), 1 mg/ml Collagenase II (Thermofisher) and 1 mg/ml Pronase K (Roche) with penicillin/streptomycin (P/S, Gibco) for 30 min at 37°C and 400 rpm. Next, isolated cells were filtered through strainer cap tubes (Falcon) and remaining red blood cells were subsequently lysed with water. The cells were stained with Annexin V-FITC/7AAD as described previously (57). Cells were acquired on an Attune NxT (Thermofisher). To determine purity of isolated IVD cells, they were washed with FACS buffer (DPBS + 5% FCS + 2mM EDTA) and stained with 1:50 pig anti-CD31 RPE (clone LCI-4) and pig anti-CD45 AF647 (K252.1E4), both BioRad.

### Transmission electron microscopy

Washed IVD punches were fixed in 2.5% Glutaraldehyde at 4°C for 72 h. Next, IVD punches were decalcified with EDTA for 10 days. After decalcification, the samples were stained and processed for image analysis as previously described (58). Images were taken with a 120 kV transmission electron microscope (FEI Tecnai G2 Spirit) equipped with two digital CCD cameras.

### Isolation and infection of human IVD cells

Human IVD tissue obtained from a total of 13 patients undergoing surgical intervention due to spinal stenosis or deformity, was fragmented mechanically and enzymatically digested overnight at 37°C and 400 rpm. Isolated cells were filtered through strainer cap tubes before seeding in tissue culture flasks (TPP) in DMEM/F12 + 10% FCS + P/S. For experiments, passage 1 cells were seeded in 96-well flat bottom plates at a density of 1×10^5^ cells per well without antibiotics. Infection at MOI 10 and intracellular survival was carried out as described previously (59).

### Assessment of IVD cell death

Cells were detached and stained with Annexin V-FITC/7AAD as previously described (56). Cells were acquired on an Attune NxT. If required, pan-caspase (50 μM Q-VD-OPH, Sigma), caspase-1 (50 μM Z-YVAD-FMK, Sigma) caspase-8 (50 μM Z-IETD-FMK, R&D Systems) or caspase-9 inhibitors (50 μM Z-LEHD-FMK, R&D Systems), respectively, were added 30 min prior to bacterial challenge. Controls were treated with DMSO only.

### Confocal laser scanning microscopy

Challenged or unchallenged IVD cells in 8-well μ slides (ibidi®) were fixed with 4% PFA. Next, they were permeabilized by Saponin and stained with 20 μM Hoechst (Themofisher) and 1.5 U Rhodamine Phalloidin (Thermofisher). Samples were visualized and acquired by CLSM with a Leica TCS SP8 inverted microscope (Leica) under a 63x/1.4 NA oil immersion objective. The obtained images were processed using Imaris 9.2.0 (Bitplane).

### Assessment of caspase activity and membrane permeability

The Caspase-Glo® 3/7, Caspase-Glo® 8 and Caspase-Glo® 9 (all Promega) kits were used as previously described (57). The Caspase-1 activity assay kit (Novus Biologicals) and the mitochondrial ToxGlo™ (Promega) assay were used accordingly to the manufacturer’s instruction. Fluorescence (488/525nm) and luminescence were measured with the SpectraMax i3 (Molecular Devices).

### Cytokine analysis

Cell culture supernatants were filtered through a 0.22 μm filter and cytokine levels were analyzed on a Luminex™ MAGPIX™ (Thermofisher) as previously described (57,59). Analysis was performed using the xPONENT® software. Data was validated additionally with the ProcartaPlex Analyst software (Thermofisher).

### Neutrophils isolation and stimulation

Neutrophils from healthy donors were isolated with the EasySep™ Direct Human Neutrophil Isolation Kit (StemCell Technologies) as previously described (57,59). Neutrophils were resuspended in DMEM/F12 and counted on an Attune NxT. They were seeded in 96-well V-well canonical plates and stimulated with either fresh medium or supernatant from un- or challenged IVD cells in a 1:3 ratio for 3 h. Neutrophils were seeded at a density of 2.5×10^5^ if not indicated differently.

### Cell surface receptor expression

Neutrophils were stained as described previously (59). Cells were stained 1:750 with the Near-IR™ Live/Dead reagent (Thermofisher), 1:50 anti-CD15 eFluor450 (clone: HI98), anti-CD181 FITC (8F1-1-4), anti-CD182 PerCP-eFluor710 (5E8-C7-F10), anti-CD183 PE-eFluor610 (CEW33D), anti-CD66b APC (G10F5), from Thermofisher and anti-CD184 BV605 (12G5) from Biolegend. Cells were acquired on an Attune NxT.

### Migration assay

For migration assays, the HTS Transwell®-96 well permeable supports with a 5 μm pore size polycarbonate membrane (Corning). In the bottom wells, either fresh medium or supernatant from un- or challenged IVD cells was placed in a 1:3 ratio. In the inserts, neutrophils were added at a density of 1×10^5^ cells. After 15 min and 45 min, the medium in the bottom well was collected and washed with FACS buffer, followed by staining with anti-CD66b for 30 min at 4°C. Absolute number of migrated neutrophils were acquired in 150 μl volume on an Attune NxT.

### ROS production

ROS production was assessed after 2.5 h of stimulation as previously described (59) with 5 μM of the CellROX™ green reagent (Thermofisher). Cells were acquired on an Attune NxT.

### Statistical analysis

Statistical analysis was done with GraphPad Prism 8. Samples were first assessed for normal distribution and then tested for statistical significance. The used tests are indicated within each figure legend.

## Acknowledgments

We’d like to thank our patients for participating in clinical and translational research, Ines Kleiber-Schaaf and Andrea Garcete for assistance with histology, Miriam Lipiski for contributing porcine spines and the center for Microscopy and Image Analysis, University of Zurich, for support with conventional, confocal laser scanning and transmission electron microscopy.

## Author contributions

TA Schweizer and AS Zinkernagel conceived the project and were involved in experimental design. TA Schweizer performed most experiments, analyzed and compiled the data. F Andreoni performed transmission electron microscopy and sample preparation. C Acevedo, TC Scheier, N Eberhard and SD Brugger collected epidemiological as well as clinical data of spondylodiscitis patients. I Heggli and S Dudli collected human intervertebral discs and provided guidance for cell isolation. E Marques Maggio performed histology. TA Schweizer and F Andreoni wrote the first draft of the manuscript. All authors helped in editing the final version of the manuscript and approved it.

