## Supplemental Material for "Intervertebral disc cell chondroptosis elicits neutrophil response in *Staphylococcus aureus* spondylodiscitis"

**Supplementary Material**

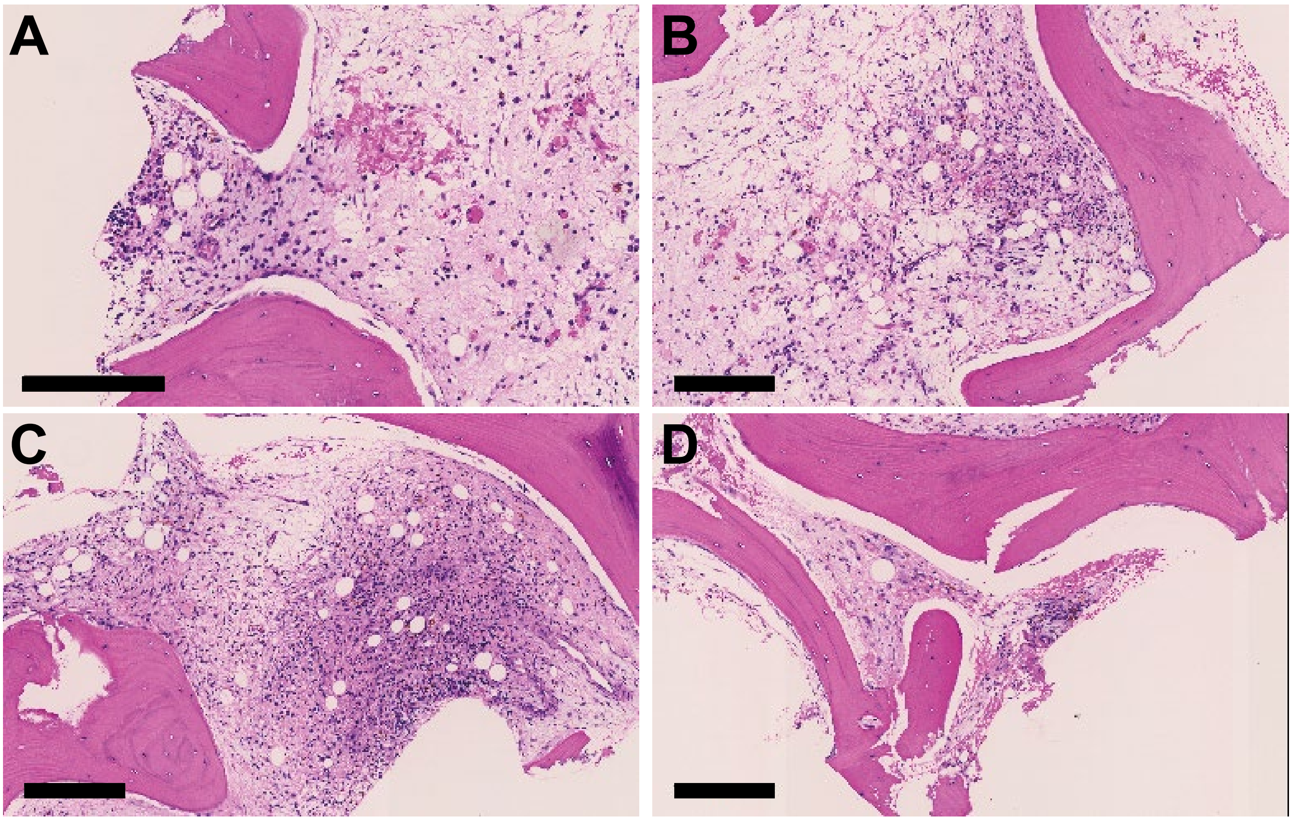


**Figure S1.** (**A-D**) H&E staining of chronic spondylodiscitis case showing presence of lymphocytes and slightly inflamed VB and IVD tissue. Scale bars indicate 200 μm. H&E, Hematoxylin & Eosin; VB, vertebral body; IVD, intervertebral disc.

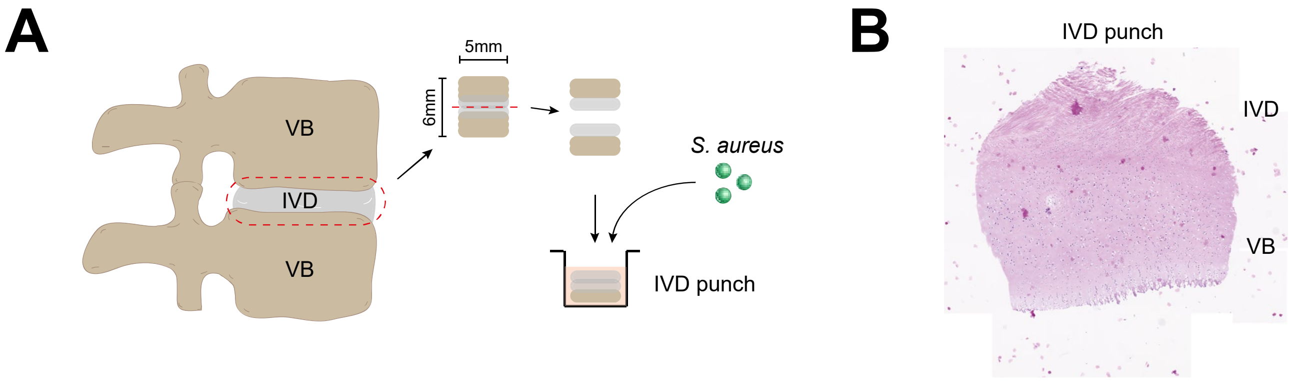


**Figure S2.** (**A**) Schematic depiction of porcine spondylodiscitis model set up. IVDs were removed from porcine spines and disc punches were prepared to be inoculated with medium only or *S. aureus* challenge. (**B**). H&E staining showing saggital section of prepared IVD punches. IVD, intervertebral disc; H&E, Hematoxylin & Eosin.

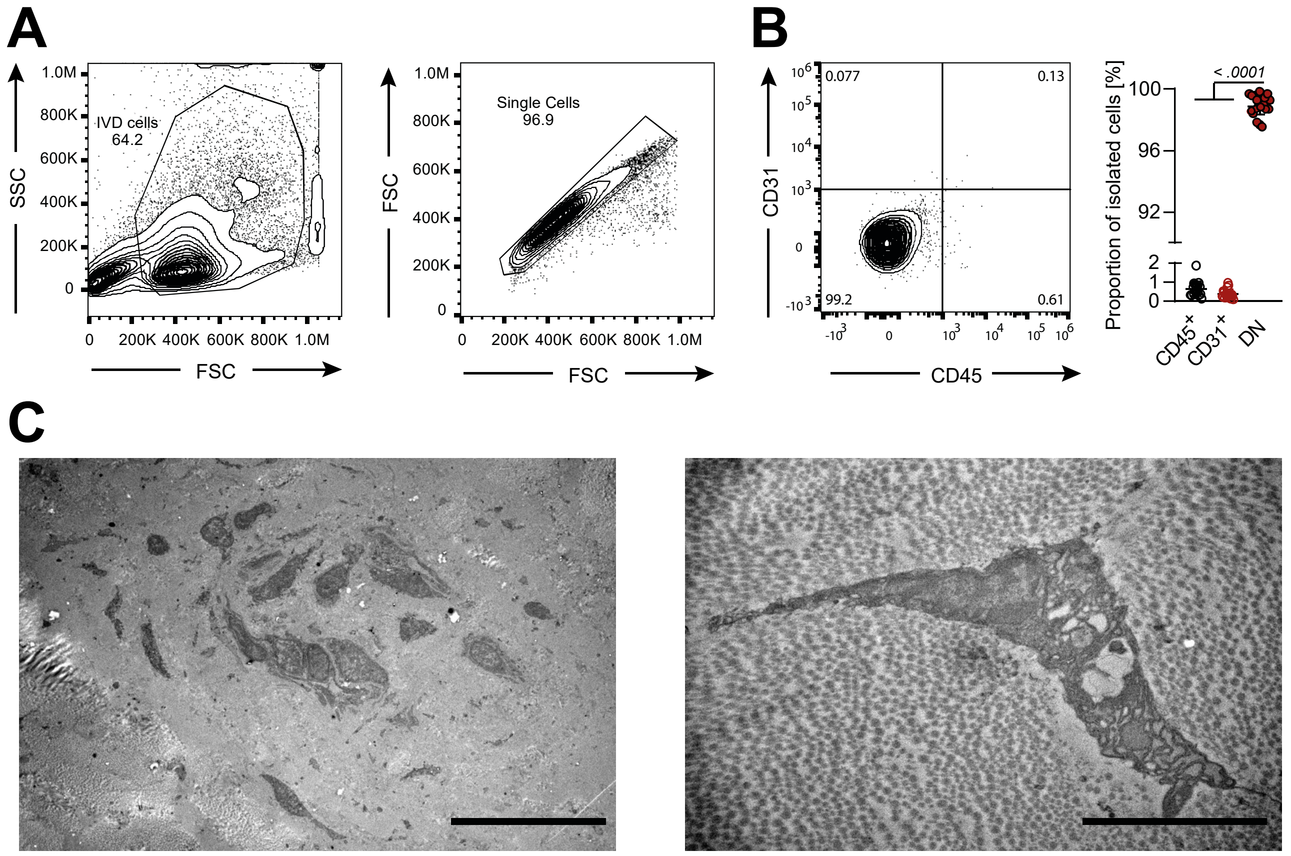


**Figure S3.** (**A**) Gating strategy of IVD cells enzymatically isolated from porcine IVD punches. (**B**) Representative flow cytometry plot and quantification of percentage CD45+, CD31+ or DN cells isolated from porcine IVD punches. (**C**) TEM of IVD cells from unchallenged porcine IVD punches. Scale bars indicate 20 µm (left panel) and 5 µm (right panel). Each dot represents isolated cells from one IVD punch. Statistical analysis was done by One-way ANOVA and Turkey's multiple comparison. IVD, intervertebral disc; DN, double negative; TEM, transmission electron microscopy.

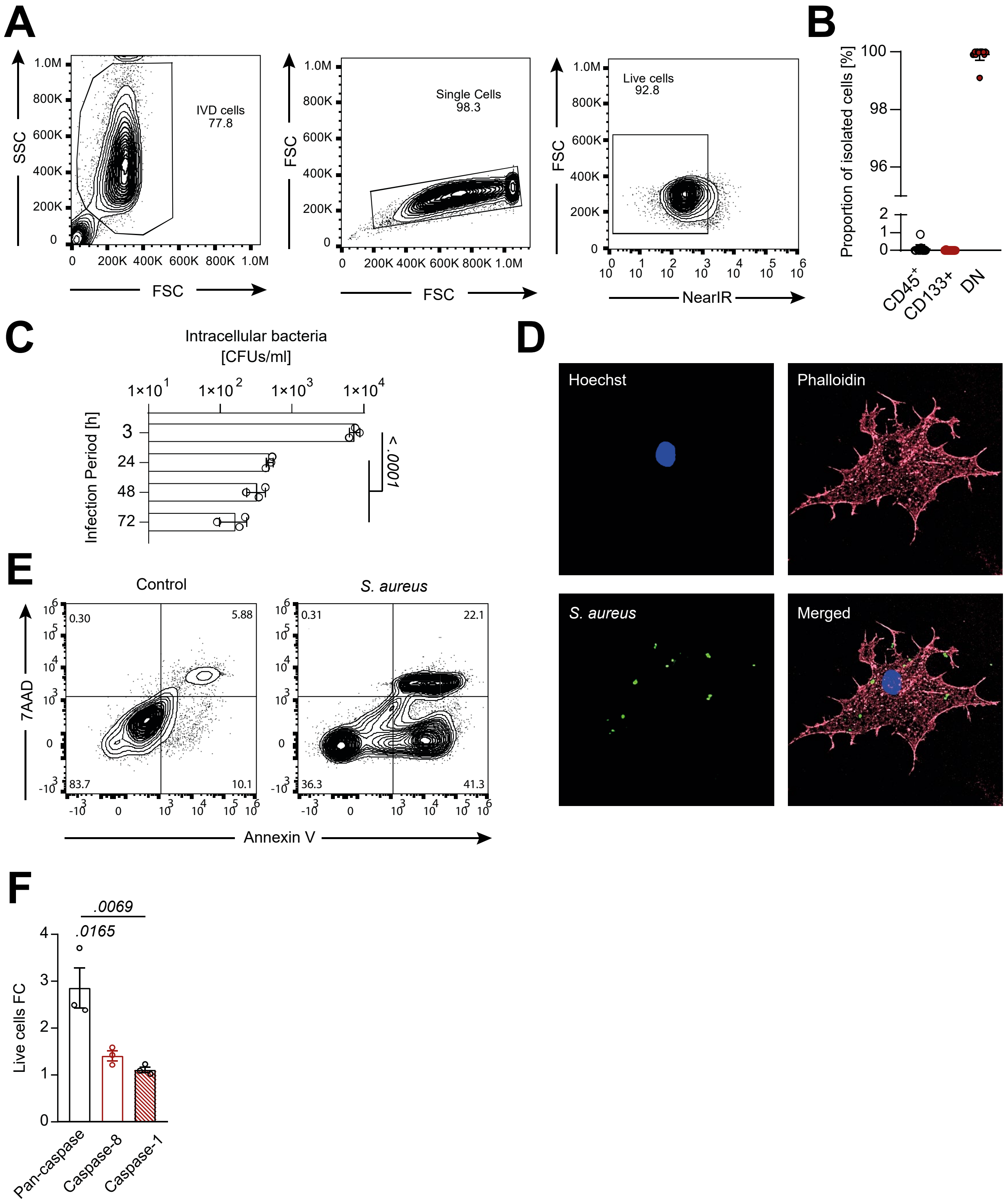


**Figure S4.** (**A and B**) Gating strategy of primary human IVD cells (A) and assessment of purity, by means of absence of CD45+ or CD133+ cells (B). (**C**) Quantification of IVD cells harboring intracellular bacteria over time. (**D**) Representative CLSM micrograph showing IVD cells harboring intracellular *S. aureus*. (**E**) Representative flow cytometry plot of cell death assessment of unchallenged or *S. aureus* challenged IVD cells. (**F**) Assessment of effect on survival of the pan-caspase (Q-VD-OP), caspase-8 (Z-IETD-FMK) and caspase-1 (Z-YVAD-FMK) inhibitors upon *S. aureus* challenge, indicated as FC to untreated IVD cells. Each dot represents one biological replicate. Statistical analysis was done by one-way ANOVA with Turkey's multiple comparison test. IVD, intervertebral disc; CLSM, confocal laser scanning microscopy; FC, fold change.

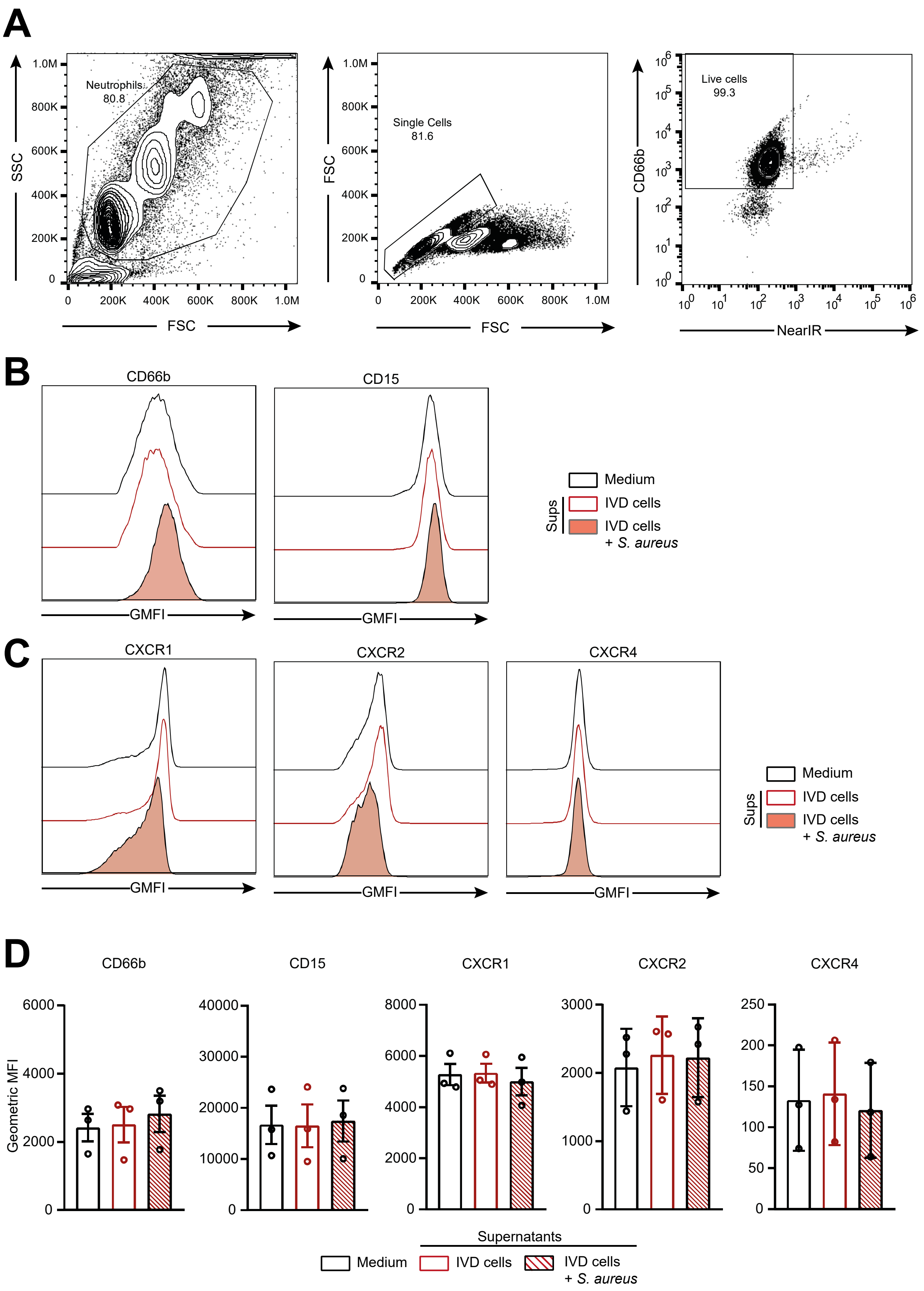


**Figure S5.** (**A**) Gating strategy of primary human neutrophils. (**B and C**) Representative flow cytometry histograms of surface CD66b and CD15 (B) as well as CXCR1, CXCR2 and CXCR4

expression (C) on neutrophils stimulated with either medium only or supernatant from 24h unchallenged or *S. aureus* challenged primary human IVD cells. (**D**) Quantification of receptor expression on neutrophils stimulated with either medium only or supernatant from 6h unchallenged or *S. aureus* challenged primary human IVD cells. IVD, intervertebral disc.

**Table S1.** ***Staphylococcus aureus*** **strains used in this study.**

| Strain | Description | Source/Reference |
|---|---|---|
| LS (JE2) | USA300 Laboratory Strain, MRSA | NARSA |
| #1 | Clinical spondylodiscitis Isolate, MSSA (Patient 6) | This study |
| #2 | Clinical spondylodiscitis isolate, MRSA (Patient 7) | This study |
| #3 | Clinical IAI spondylodiscitis isolate, MRSA | This study |

LS, laboratory strain; MRSA, Methicillin-resistant *S. aureus*; NARSA, Network on Antimicrobial Resistance in *S. aureus*; MSSA, Methicillin-susceptible *S. aureus*; IAI, Implant-associated infection.
